# A Controlled Slip: Training Propulsion via Acceleration of the Trailing Limb

**DOI:** 10.1101/2020.03.27.012591

**Authors:** Andria J. Farrens, Maria Lilley, Fabrizio Sergi

## Abstract

Walking function, which is critical to performing many activities of daily living, is commonly assessed by walking speed. Walking speed is dependent on propulsion, which is governed by ankle moment and the posture of the trailing limb during push-off. Here, we present a new gait training paradigm that utilizes a dual belt treadmill to train both components of propulsion by accelerating the belt of the trailing limb during push off. Accelerations require subjects to produce greater propulsive force to counteract inertial effects, and increase trailing limb angle through increased belt velocity.

We hypothesized that exposure to our training p rogram would produce after effects in propulsion mechanics and, consequently, walking speed. We tested our protocol on healthy subjects at two acceleration magnitudes–Perceptible (PE), and Imperceptible, (IM)–and compared their results to a third control group (VC) that walked at a higher velocity during training.

Results show that the PE group significantly increased walking speed following training (mean ± s.e.m: 0.073 ± 0.013 m/s, p < 0.001). The change in walking speed in the IM and VC groups was not significant at the group level (IM: 0.032 ± 0.013 m/s; VC: -0.003 ± 0.013 m/s). Responder analysis showed that changes in push-off posture and in neuro-motor control of ankle plantar-flexor muscles contributed to the larger increases in gait speed measured in the PE group compared to the IM and VC groups. Analysis of the effects during and after training suggest that changes in neuromotor coordination are consistent with use-dependent learning.

## I. Introduction

Ambulation is critical to performing activities of daily living, including self-care and community engagement. Walking ability, commonly assessed by walking speed, decreases with age and is affected by numerous neurological conditions [1], [2]. As our aging population increases, there is a critical need for rehabilitation techniques that are effective in retraining walking ability to prolong independent living and quality of life for these individuals.

Walking function is dependent on propulsive force generation in both young and elderly individuals [1], [3]. Propulsive force generation is determined by two factors: posture of the trailing limb at push-off and ankle moment [2], [4]–[6]. Posture that increases the distance between the foot of the trailing limb and body center of mass at push off increases propulsion by increasing the component of the anterior ground reaction force that acts in the forward direction. These postural changes can be quantified by trailing limb angle and stride length. While postural modification alone can increase propulsion, changes in ankle plantarflexor muscles that generate ankle moment also contribute to propulsion. Moreover, decreases in ankle moment have been highlighted as a functional limitation in elderly adults that leads to decreased walking speed, making it a key target for intervention [3].

In this work, we present a novel paradigm for goal-oriented gait training that utilizes a dual belt treadmill to train both components of propulsion, i.e. push-off posture and ankle moment. This is achieved by accelerating the treadmill belt of the trailing limb during the double support phase of walking. The belt acceleration introduces a fictitious inertial force that requires the ankle plantarflexors to apply greater forces during push-off to maintain a steady position on the treadmill. Assuming no modification in push-off timing, accelerations of the belt cause the trailing limb to be moved at a larger velocity that results in an increase in trailing limb angle and stride length, training modifications in push off posture that increase propulsion through mechanical advantage.

Behavioral after effects of our training paradigm could be driven by two independent learning processes: adaptation or use-dependent learning (UDL). Adaptation is a learned response to a change in environmental dynamics that drives a recalibration of feedforward motor commands that persist when the environmental demands are removed [11], [12]. Larger (perceptible) environmental perturbations are thought to stimulate more explicit motor control centers that drive fast reactive changes and after effects that decay quickly, while smaller (imperceptible) perturbations may stimulate slower, implicit motor learning that results in longer lasting retention of the adaptated behavior [13]. Locomotor adaptation approaches have been used in gait rehabilitation to address gait asymmetry [12] and foot clearance [14], but have not been used to directly affect propulsion. Instead, UDL is a type of Hebbian learning that is the basis of many high-repetition rehabilitation protocols [15]–[17]. UDL occurs when subjects perform numerous consistent movements that bias future movements in the direction of the repeated movement. In our protocol, repeated movements consist of a more mechanically advantagous push-off posture and increased ankle moment torque.

The primary purpose of this study was to test the efficacy of training propulsion using belt accelerations. We tested our paradigm in healthy individuals at two acceleration magnitudes (Perceptible, PE and Imperceptible, IM), and included a velocity control group (VC) that walked at a higher velocity during training rather than experiencing belt accelerations. The VC group was matched to the velocity change imposed in the PE acceleration training group. To test for effects in our primary outcome measure, gait speed, we evaluated subjects self-selected treadmill walking speed before and after exposure to training in a user driven speed condition [18]. To quantify modifications in push of posture, stride length and trailing limb angle (TLA) were measured via 3D motion tracking. Modifications in ankle moment were quantified via EMG data measured from four ankle plantar-or dorsi-flexor muscles and from force-plate measurements of the anterior ground reaction force (AGRF) during push off.

We hypothesized that exposure to our training paradigm would produce after effects that increase walking speed in both acceleration training groups, but not in the velocity control group. Measured changes in walking speed may be driven by changes in propulsive force generation, due to changes in walking kinematics and/or kinetics. We hypothesized that there would be an increase in TLA, SL, propulsive impulse, peak AGRF, and plantar-flexor activation in all groups during training, but sustained change following training only in the acceleration training groups. To determine if after effects of training were driven by adaptation or UDL, we evaluated the evolution of effects in each gait parameter during and after training to determine which type of learning best described measured patterns of behavior.

## II. Materials and Methods

A total of 76 healthy individuals participated in this study. Of the 76 individuals, eight were excluded due to technical failure, four for self-selected walking speed outside the range [0.6 1.6] m/s, two for excessive cross-over between treadmill belts during walking, two for highly variable walking speed during the user-driven treadmill conditions (> ±0.05 m/s within a minute), and one for tripping. The Perceptible (PE) group had nineteen subjects (9 males, age (mean ± std) 24 ± 4 y), the Imperceptible (IM) group had twenty subjects (10 males, age 23.5 ± 3.75 y), and the Velocity Control (VC) group had twenty subjects (11 males, age 24.5 ± 3.5 y). The study was approved by the Institutional Review Board of the University of Delaware (IRBNet ID: 929630-5) and each subject provided written informed consent and receive compensation for participation.

### A. Experimental Set-Up

Our experimental protocol is shown in Fig. 1. Participants walked for 5 minutes at a self-selected speed (Baseline 1), followed by 10 minutes in the training condition, and a final 5 minutes at a self-selected speed (Baseline 2). In Baseline 1 and 2 the treadmill speed was set via a user-driven speed controller described in Sec. II-C. Subjects walked on an instrumented dual-belt treadmill (Bertec Corp., Columbus OH, USA), while wearing four reflective spherical markers (two per greater trochanter, two per lateral malleolus), and sixteen bipolar EMG electrodes (bilaterally on the tibialis anterior, lateral gastrocnemius, medial gastrocnemius, and soleus muscles). A ten camera Vicon T40-S passive motion capture system (Oxford Metrics, Oxford, UK) was used to measure marker position in 3D space and an OT Bioelettronica amplifier and software were used to acquire EMG. Due to system limitations, marker data were acquired only during the four periods highlighted in Fig. 1, at 100 Hz. EMG and treadmill analog force/torque data were acquired continuously throughout all experimental conditions at 10,240 Hz and 500 Hz respectively. A 24-in screen in front of the treadmill provided feedback on protocol duration and a visual target to keep subjects from looking down at the treadmill. Subjects wore noise cancelling headphones (COWIN E7) that played white noise to eliminate environmental distractions.

**Fig. 1.**
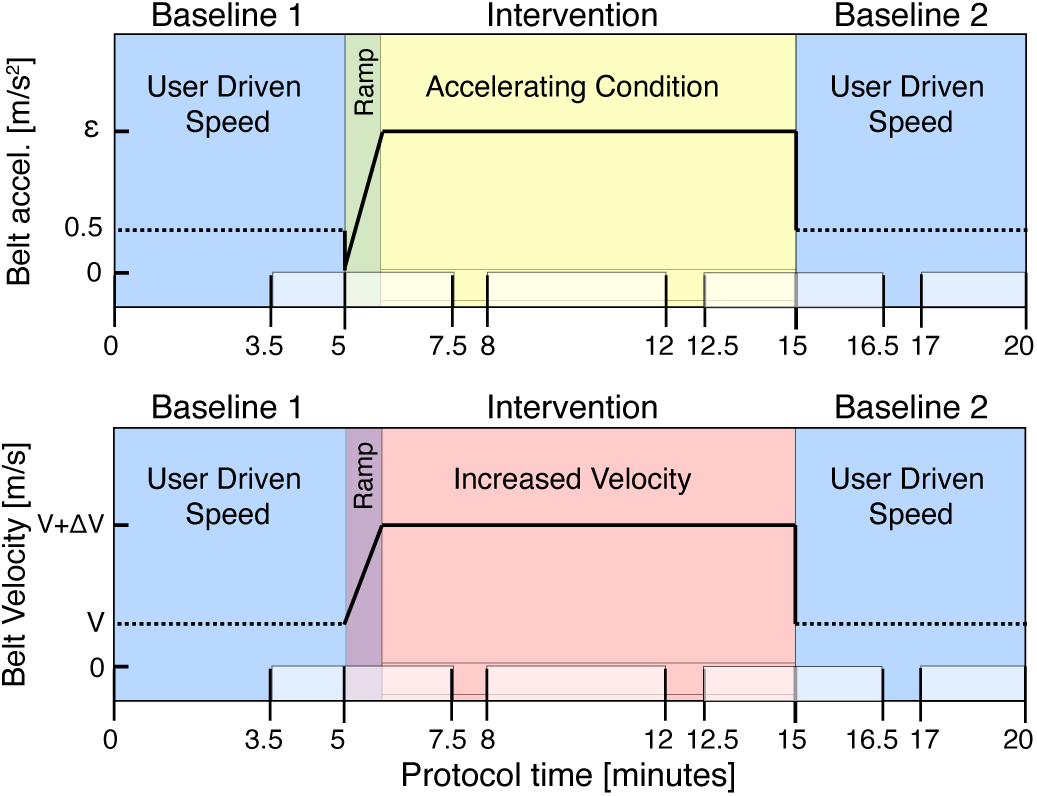
Experimental Protocol Schematic. For both Baseline conditions, subjects walked in the UDTC, with maximum belt accelerations capped at 0.5 m/s^2^. Highlighted phases at the bottom signify periods in which kinematic marker data were collected. Transition to all training conditions included a minute long ramp phase to gradually introduce belt accelerations or change in velocity. Top: Accelerating intervention. Belt accelerations are shown on the y-axis, where *ϵ* signifies the magnitude of accelerations applied during intervention (either 2 m/s^2^ or 7 m/s^2^). Bottom: Velocity Control Intervention. Belt velocity is shown on the y-axis, an increase of *δ*V = 0.05 m/s was applied to the end velocity of Baseline 1.

The perceptible acceleration magnitude was set to 7 m/s^2^, which was determined to be highly noticeable and safe for healthy individuals. An imperceptible acceleration threshold of 2 m/s^2^ was established via testing on a separate set of 10 healthy individuals. For the VC group, an increase in velocity of 0.05 m/s over subjects baseline walking speed was imposed. The change in velocity was chosen to match the change in velocity experienced in the PE group, determined from 9 subjects from our previous study [19]. Data from eight subjects in the imperceptible training group in our previous study were included in IM group in this study.

### B. Self-selected walking speed

A preliminary set of trials were conducted to determine subjects self-selected walking speed (SS-WS) immediately prior to our protocol. First, subjects fastest comfortable walking speed (FC-WS) was ascertained. Participants walked on the treadmill at an initial speed of 0.5 m/s that was increased by the experimenter in intervals of 0.02 m/s until the participant verbally indicated the treadmill had reached their FC-WS. The treadmill was returned to 0.5 m/s and the same ramp-up procedure was then repeated to find subjects SS-WS. A ramp-down procedure was then conducted with the treadmill starting at the subjects FC-WS and decreased in increments of 0.02 m/s until the subject indicated their SS-WS had been reached. The ramp-up and ramp-down procedures to determine SS-WS were each repeated twice, and the average of the four measured velocities was taken as the subjects SS-WS [10].

### C. User-Driven Speed Controller

In standard treadmill walking, walking speed is restricted to the constant velocity imposed by the treadmill. Because our training paradigm seeks to modify subjects walking speed, a treadmill that operates at a constant velocity–and thus restricts changes in walking speed–is impractical, and is likely to eliminate any after effects acceleration training may induce. To address this, we used a user-driven treadmill controller (UDTC) that changes the velocity of the treadmill in response to changes in the subjects walking behavior, and more readily mirrors over-ground walking conditions [18].

The UDTC changes speed based on an empirically weighted combination of the following three gait parameters: change in AGRF, step length, and center of mass position relative to the center of the treadmill. For example, if subjects produced greater AGRF, increased their step length, or walked further forward on the treadmill, the treadmill speed would increase. Conversely, decreases in AGRF, step length, or movement to the back of the treadmill would decrease speed. The maximum belt acceleration was set to 0.5 m/*s*^2^, and was previously tested to ensure subject safety and comfort [18], [19].

Prior to our experimental protocol, participants were given a brief training in the UDTC condition. Participants were started at their SS-WS and given up to 5 minutes to familiarize themselves with the UDTC. In line with previous results, several subjects increased their walking speed on the UDTC from their initial SS-WS chosen in the fixed speed condition [18]. Consequently, for our experimental protocol, the initial baseline period of walking was set to at least 5 minutes, but was continued until peak change in velocity within one minute was smaller than 0.05 m/s. The Baseline 1 (Fig. 1) condition of our protocol was then defined as the period spanning the one minute of steady state walking, and the previous four minutes prior to acheiving a steady state.

### D. Belt Acceleration Controller

To train increases in propulsive force generation, we developed a controller capable of accelerating the treadmill belt of the trailing limb during the double support phase of gait and returning it to a nominal speed during the swing phase of gait (Fig. 2). The rationale behind the selection of this dynamic distortion (acceleration) was to attenuate subjects’ propulsive force by introducing a fictitious inertial force in the opposite direction. Moreover, assuming that subjects do not modify their push-off timing, the foot on the accelerated belt will move at a larger average speed, causing TLA and SL to increase. In this way, we aimed to target both gait mechanisms (ankle moment and posture of the trailing limb at push-off) that control propulsive force generation.

**Fig. 2.**
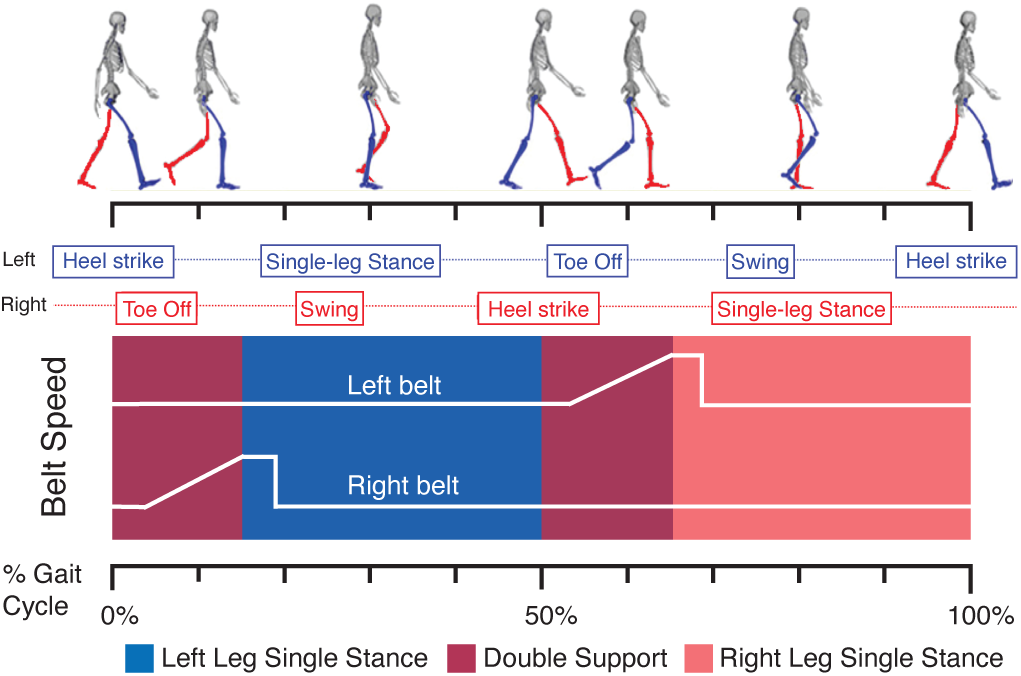
Belt acceleration during intervention as a function of gait cycle

Push off occurs at the end of the double support phase of gait that typically lasts for 100-150 ms. While double support can be detected in real time using force-plate data, the delay between detection of dual support and the execution of an acceleration command exceeded 150 ms, making the use of real-time detection impracticable. As such, we developed a simple algorithm to predict when push off would occur based on the prior gait cycle that included an anticipation factor to account for our system delays. The controller sends an acceleration signal at a time *t*, defined as the instant at which the condition below becomes true

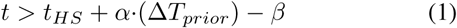

where *t* is the current time, *t*_*HS*_ is the time instant of heel strike of the leg currently in stance, Δ*T*_*prior*_ is the time between the previous left and right heel strikes that provides a prediction of when double support will occur, and *β* is the anticipation factor. Based on this logic, when the amount of time elapsed following heel strike exceeds *α* (Δ*T*_*prior*_) − *β*, the acceleration signal is sent. Parameter values of *α* = 1.175 and *β* = 0.185 s were determined empirically on a separate group of 10 individuals to confirm accelerations occured during pushoff at a reasonable range of speeds (0.7 - 1.4 m/s). Accelerations were saturated by a speed increase limit of 0.7 m/s to ensure subjects safety. 100 ms after detection of toe-off, the controller decelerates the belt back to the nominal speed during the swing phase (Fig. 2).

#### 1) Data Preprocessing

EMG, kinematic marker data, and gait speed data were acquired on 3 separate systems and time synced via a common force-plate data signal. VICON marker position data were fed into a standard Visual3D preprocessing pipeline, which included i) manual labelling of markers; ii) interpolation of missing marker data with a third order polynomial fit for a maximum gap size of five samples; and iii) low-pass filtering at 6 Hz with a 4th order zero-shift Butterworth filter [10]. Force-plate data were filtered with a 4th order zero-shift low-pass Butterworth filter at 25 Hz [10]. EMG data were bandpass filtered at 20-500 Hz, rectified, and the envelope was taken via a lowpass zero-shift 4th order Butterworth filter with a 10 Hz cut off frequency [5].

#### 2) Data Analysis

Force-plate data were used to define heel strike and toe off events. Heel strike events were determined as the instants at which the vertical ground reaction force exceeded 5% max force and remained above 5% max force for at least 200 ms. Toe off events were determined as the instants at which the vertical ground reaction force fell below 5% max force and remained below for at least 150 ms. EMG data of each muscle (tibialis anterior, lateral and medial gastrocnemius, and soleus muscles) were segmented and linearly resampled to [0 - 100] percent of gait cycle, defined by each heel strike to heel strike event. Each gait cycle was further divided into periods of single and double support using heel-strike and toe-off events (Fig. 4). To quantify peak EMG activation for plantar-flexor muscles related to propulsion, we took the peak EMG signal measured during the single support phase (12-50% GC) [5], [6]. To quantify peak EMG activation for dorsiflexor muscles related to weight acceptance, we calculated the peak EMG signal in the interval around heel strike (0-12, 80-100% GC) [5], [6]. EMG cabling noise introduced by walking excluded the use of peak EMG measured in one subject in each of the acceleration training groups. EMG data were normalized by the median peak activation measured during the last minute of baseline 1 walking when subjects had reached a steady state velocity.

**Fig. 3.**
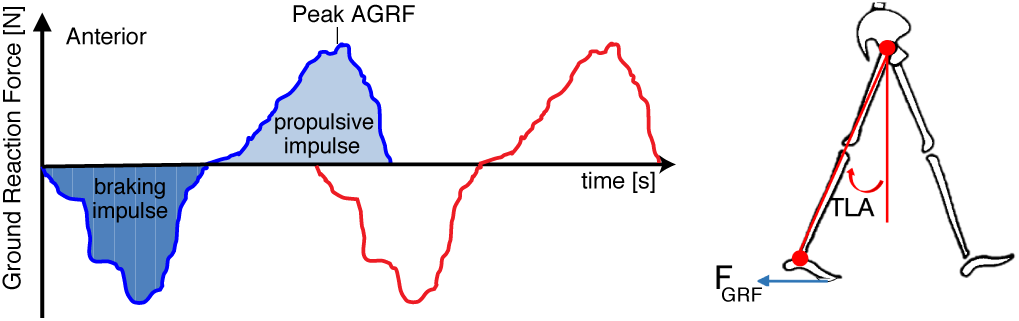
Definition of gait parameters propulsive impulse (PI) and Peak AGRF (left) and TLA (right)

**Fig. 4.**
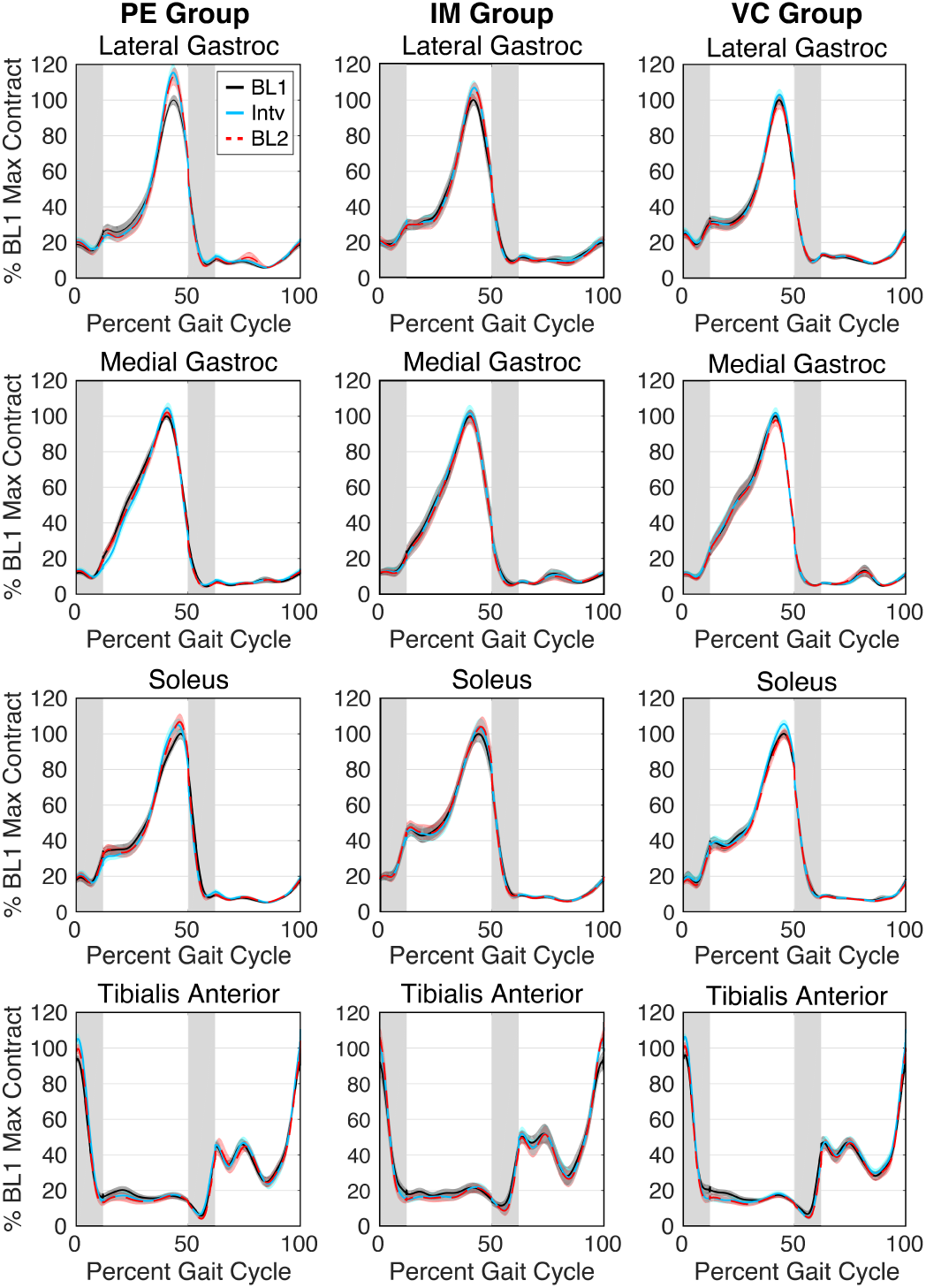
Group average plantar flexor and dorsiflexor EMG activation, resampled over gait cycle (HS-HS). Grey pannels denote double support phases. The average activation for each subject in each experimental phase— BL1, Intv. (Early and Late), BL2 (Early and Late)—was used to calculate the group average (solid/dashed lines) and standard error (shaded area).

Propulsive impulse (PI) was calculated from the force-plate data as the area under the positive (anterior) portion of the ground reaction force (Fig. 3) [2]. Peak AGRF was taken as the maximum anterior ground reaction force. Stride length was calculated as the anterior-posterior deviation of the lateral malleolus ankle marker from heel strike to heel strike [20]. TLA was defined as the maximum angle between the straight line connecting the greater trochanter and the lateral malleolus of the trailing limb for each stride cycle (Fig. 3). TLA is typically taken at time of peak AGRF (not max angle) [2], however distortions in force-plate measures due to belt acceleration invalidated this measure during both accelerating training conditions [21]. Our alternative max TLA measure was highly correlated (*R*^2^ = 0.98) with TLA measured at max AGRF in BL1 for all groups, and is not a large departure from the traditional measure, and has been used before in quantifying paretic TLA [22]. Gait speed (GS) was sampled at right and left heelstrike events, as the UDTC is designed to update belt speed at each heel strike event.

### E. Statistical Analysis

We used a mixed model analysis to evaluate the effects of training and training type on our gait parameters (GS, SL, TLA, Peak EMG, PI, and Peak AGRF). Training type was set as the between subjects factor (PE, IM, VC). Within-subjects factors included baseline 1 (BL1), Early and Late Intervention (Early Intv., Late Intv), and Early and Late baseline 2 (Early BL2, Late BL2). BL1 was calculated as the mean of the last minute of baseline 1 walking. Early and Late Intv. were calculated as the mean values measured in the first and last 20 strides in the training condition (not including the ramp phase), and were include to determine the **effects of training** for all valid gait measures (TLA, SL, and EMG). Early and Late BL2 were calculated as the mean values measured in strides 5-24 and last 20 strides in the baseline 2 condition, and were included to determine the **after effects** of training on all gait parameters. Baseline 2 strides 1-4 were excluded due to transient effects of subjects taking “stutter steps” as a result of the treadmill behavior changing midstream (see supplemental) [20]. When the mixed model returned a significant effect, Tukey HSD post-hoc testing was used to quantify the effect of training on the measured gait parameters.

To investigate efficacy of training at the individual level, we calculated z-score of the within subject change from BL1 in the experimental phases Early and Late BL2, defined as before. We then classified subjects as positive responders if their z-score was greater than zero, and negative responders if their z-score was less than zero, and quantified the number of positive and negative responders in each group following training. For all outcome measures, the mean was obtained by averaging right and left leg data. MATLAB and JMP Pro software was used for all statistical analyses.

## III. Results

### A. Gait Speed

The model returned a significant fixed effect of experimental phase, and an interaction between training type and experimental phase (Tbl. I). No significant effect of group was detected. Model parameter estimates showed that across groups late BL2 was significantly greater than both BL1 and Early BL2 (mean ± s.e.m: 0.034 ± 0.007 m/s, *p* < 0.001; 0.023 ± 0.007 m/s, *p* = 0.008, respectively). Model parameter estimates showed that the interaction was driven by a greater increase from Early to Late BL2 in the PE group compared the to VC group (**PE:** 0.052 ± 0.013 m/s; **VC:** -0.009 ± 0.013 m/s, *p* = 0.001), and a greater increase in Late BL2 compared to BL1 in the PE group compared to both the IM and VC groups (**PE:** 0.073 ± 0.013 m/s, **IM:** 0.032 ± 0.013 m/s; **VC:** -0.003 ± 0.013 m/s; **PE vs IM:** *p* = 0.031, **PE vs VC:** *p* < 0.001). Post-hoc Tukey HSD testing revealed a significant increase in gait speed in Late BL2 compared to BL1 and Early BL2 in the PE group (0.073 ± 0.013 m/s, *p* < 0.001; 0.052 ± 0.013 m/s, *p* = 0.005, respectively). No significant change in gait speed was detected in either the IM or VC groups.

**TABLE I.** Mixed Model Results: Fixed Effects for all gait parameters.

| Mixed Model: Fixed Effects Tests |  |  |  |  |  |
| --- | --- | --- | --- | --- | --- |
| Gait Speed | Nparm | DF Num. | DF Den. | F Ratio | Prob > F |
| Group | 2 | 2 | 59.80 | 0.608 | 0.548 |
| Exp. Phase | 2 | 2 | 112.00 | 10.552 | < 0.001 |
| Group*Exp. Phase | 4 | 4 | 112.00 | 4.621 | 0.002 |
| TLA | Nparm | DF Num. | DF Den. | F Ratio | Prob > F |
| Group | 2 | 2 | 59.00 | 0.726 | 0.488 |
| Exp. Phase | 4 | 4 | 223.00 | 20.583 | < 0.001 |
| Group*Exp. Phase | 8 | 8 | 223.00 | 1.795 | 0.079 |
| Stride Length | Nparm | DF Num. | DF Den. | F Ratio | Prob > F |
| Group | 2 | 2 | 61.20 | 1.299 | 0.280 |
| Exp. Phase | 4 | 4 | 223.00 | 10.192 | < 0.001 |
| Group*Exp. Phase | 8 | 8 | 223.00 | 1.863 | 0.067 |
| Prop. Impulse | Nparm | DF Num. | DF Den. | F Ratio | Prob > F |
| Group | 2 | 2 | 58.50 | 0.12 | 0.890 |
| Exp. Phase | 2 | 2 | 112.00 | 34.189 | < 0.001 |
| Group*Exp. Phase | 4 | 4 | 112.00 | 2.091 | 0.087 |
| Peak AGRF | Nparm | DF Num. | DF Den. | F Ratio | Prob > F |
| Group | 2 | 2 | 59.10 | 0.292 | 0.748 |
| Exp. Phase | 2 | 2 | 112.00 | 18.603 | < 0.001 |
| Group*Exp. Phase | 4 | 4 | 112.00 | 4.177 | 0.003 |
| Lateral Gast. | Nparm | DF Num. | DF Den. | F Ratio | Prob > F |
| Group | 2 | 2 | 203.30 | 0.000 | 1.000 |
| Exp. Phase | 4 | 4 | 216.00 | 15.480 | < 0.001 |
| Group*Exp. Phase | 8 | 8 | 216.00 | 2.775 | 0.006 |
| Medial Gast. | Nparm | DF Num. | DF Den. | F Ratio | Prob > F |
| Group | 2 | 2 | 213.40 | 0.000 | 1.000 |
| Exp. Phase | 4 | 4 | 216.00 | 11.269 | < 0.001 |
| Group*Exp. Phase | 8 | 8 | 216.00 | 0.855 | 0.555 |
| Soleus | Nparm | DF Num. | DF Den. | F Ratio | Prob > F |
| Group | 2 | 2 | 190.40 | 0.000 | 1.000 |
| Exp. Phase | 4 | 4 | 216.00 | 11.691 | < 0.001 |
| Group*Exp. Phase | 8 | 8 | 216.00 | 2.434 | 0.015 |
| Tibialis Ant. | Nparm | DF Num. | DF Den. | F Ratio | Prob > F |
| Group | 2 | 2 | 232.70 | 0.000 | 1.000 |
| Exp. Phase | 4 | 4 | 216.00 | 8.301 | < 0.001 |
| Group*Exp. Phase | 8 | 8 | 216.00 | 2.937 | 0.004 |

Responder analysis (Tbl. II, Fig. 5) showed a greater number of positive responders and larger median effects in the PE group across BL2, and in the IM group in late BL2 (**PE:** Early: 14 positive responders (median z-score: 2.39, range: [0.59 6.25]), Late: 15 (8.36 [1.36 20.91]); **IM:** Early: 9 (1.74 [0.56 8.69]), Late: 14 (4.48 [0.03 25.52]); **VC:** Early: 14 (1.22 [0.01 3.05]), Late: 10 (3.09 [0.08 16.56])).

**TABLE II.** Responder Analysis: median effect size and range.

|  | Perceptible |  | Imperceptible |  | Velocity Control |  |
| --- | --- | --- | --- | --- | --- | --- |
| GS | Num | Median [range] | Num | Median [range] | Num | Median [range] |
| Early Pos | 14 | 2.39 [0.59 6.25] | 9 | 1.74 [0.56 8.69] | 14 | 1.22 [0.01 3.05] |
| Early Neg | 5 | -0.83 [-2.31 -0.10] | 11 | -0.72 [-5.29 -0.01] | 6 | -1.08 [-1.93 -0.14] |
| Late Pos | 15 | 8.36 [1.36 20.91] | 14 | 4.48 [0.03 25.52] | 10 | 3.09 [0.08 16.56] |
| Late Neg | 4 | -4.40 [-6.88 -1.63] | 6 | -2.76 [-7.08 -0.81] | 10 | -4.15 [-7.12 -2.15] |
| TLA | Num | Median [range] | Num | Median [range] | Num | Median [range] |
| Early Pos | 16 | 1.27 [0.13 2.80] | 14 | 1.42 [0.06 3.41] | 11 | 1.08 [0.16 2.24] |
| Early Neg | 3 | -0.69 [-0.78 -0.65] | 6 | -0.20 [-1.15 -0.06] | 9 | -0.64 [-3.68 -0.09] |
| Late Pos | 14 | 2.24 [0.32 5.03] | 15 | 1.14 [0.05 5.08] | 12 | 1.00 [0.04 2.30] |
| Late Neg | 5 | -0.80 [-1.39 -0.15] | 5 | -0.83 [-2.07 -0.08] | 8 | -0.86 [-1.50 -0.45] |
| SL | Num | Median [range] | Num | Median [range] | Num | Median [range] |
| Early Pos | 17 | 1.23 [0.24 3.36] | 14 | 1.38 [0.05 5.09] | 11 | 0.72 [0.07 2.47] |
| Early Neg | 2 | -0.92 [-1.56 -0.29] | 6 | -0.16 [-1.32 -0.01] | 9 | -0.45 [-3.24 -0.14] |
| Late Pos | 13 | 2.78 [1.13 5.37] | 11 | 2.68 [0.27 9.53] | 10 | 1.16 [0.46 2.73] |
| Late Neg | 6 | -0.63 [-1.12 -0.07] | 9 | -0.73 [-2.12 -0.07] | 10 | -0.77 [-1.64 -0.07] |
| PI | Num | Median [range] | Num | Median [range] | Num | Median [range] |
| Early Pos | 14 | 0.69 [0.06 2.36] | 15 | 0.98 [0.07 2.99] | 15 | 0.83 [0.10 3.72] |
| Early Neg | 5 | -0.69 [-1.75 -0.31] | 5 | -0.21 [-0.42 -0.01] | 5 | -0.72 [-1.07 -0.12] |
| Late Pos | 16 | 2.06 [0.31 4.46] | 18 | 0.83 [0.01 5.48] | 17 | 0.75 [0.23 3.34] |
| Late Neg | 3 | -0.11 [-1.10 -0.00] | 2 | -0.37 [-0.52 -0.22] | 3 | -0.25 [-0.73 -0.12] |
| P. AGRF | Num | Median [range] | Num | Median [range] | Num | Median [range] |
| Early Pos | 11 | 1.13 [0.00 1.62] | 13 | 0.92 [0.04 2.34] | 17 | 0.75 [0.06 2.29] |
| Early Neg | 8 | -0.61 [-1.91 -0.04] | 7 | -0.49 [-0.87 -0.20] | 3 | -0.82 [-0.94 -0.55] |
| Late Pos | 14 | 2.98 [0.66 5.35] | 17 | 0.72 [0.12 5.65] | 12 | 1.23 [0.06 2.55] |
| Late Neg | 5 | -0.87 [-1.53 -0.10] | 3 | -1.18 [-1.71 -0.58] | 8 | -0.66 [-1.48 -0.05] |
| LG | Num | Median [range] | Num | Median [range] | Num | Median [range] |
| Early Pos | 13 | 0.74 [0.10 1.70] | 10 | 0.39 [0.09 2.23] | 14 | 0.40 [0.02 1.69] |
| Early Neg | 5 | -0.44 [-0.53 -0.11] | 9 | -0.27 [-0.65 -0.05] | 6 | -0.30 [-0.91 -0.11] |
| Late Pos | 14 | 0.93 [0.02 3.75] | 10 | 0.55 [0.20 3.03] | 11 | 0.15 [0.01 1.25] |
| Late Neg | 5 | -0.46 [-0.76 -0.23] | 9 | -0.30 [-0.83 -0.03] | 9 | -0.42 [-0.87 -0.04] |
| MG | Num | Median [range] | Num | Median [range] | Num | Median [range] |
| Early Pos | 9 | 0.36 [0.03 0.89] | 9 | 0.18 [0.01 0.82] | 12 | 0.22 [0.01 0.72] |
| Early Neg | 9 | -0.42 [-0.96 -0.07] | 10 | -0.58 [-0.75 -0.11] | 8 | -0.33 [-0.84 -0.11] |
| Late Pos | 9 | 0.30 [0.01 1.65] | 8 | 0.22 [0.02 1.15] | 9 | 0.36 [0.03 1.01] |
| Late Neg | 9 | -0.23 [-0.69 -0.02] | 11 | -0.34 [-1.29 -0.02] | 11 | -0.37 [-1.35 -0.10] |
| SO | Num | Median [range] | Num | Median [range] | Num | Median [range] |
| Early Pos | 14 | 0.50 [0.06 1.38] | 12 | 0.30 [0.09 1.46] | 7 | 0.57 [0.05 1.20] |
| Early Neg | 4 | -0.06 [-0.74 -0.01] | 7 | -0.15 [-0.70 -0.06] | 13 | -0.23 [-0.84 -0.01] |
| Late Pos | 14 | 0.63 [0.10 2.95] | 11 | 0.88 [0.15 2.82] | 7 | 0.34 [0.05 2.17] |
| Late Neg | 4 | -0.14 [-1.45 -0.03] | 8 | -0.05 [-0.72 -0.00] | 13 | -0.29 [-1.13 -0.02] |
| TA | Num | Median [range] | Num | Median [range] | Num | Median [range] |
| Early Pos | 11 | 1.00 [0.27 1.21] | 11 | 0.26 [0.10 0.93] | 12 | 0.31 [0.11 1.24] |
| Early Neg | 7 | -0.33 [-1.70 -0.12] | 8 | -0.18 [-0.89 -0.03] | 8 | -0.24 [-0.78 -0.02] |
| Late Pos | 12 | 0.97 [0.08 2.82] | 12 | 0.45 [0.12 2.22] | 11 | 0.27 [0.04 1.56] |
| Late Neg | 6 | -0.22 [-0.87 -0.01] | 7 | -0.35 [-1.47 -0.02] | 9 | -0.26 [-1.16 -0.05] |

**Fig. 5.**
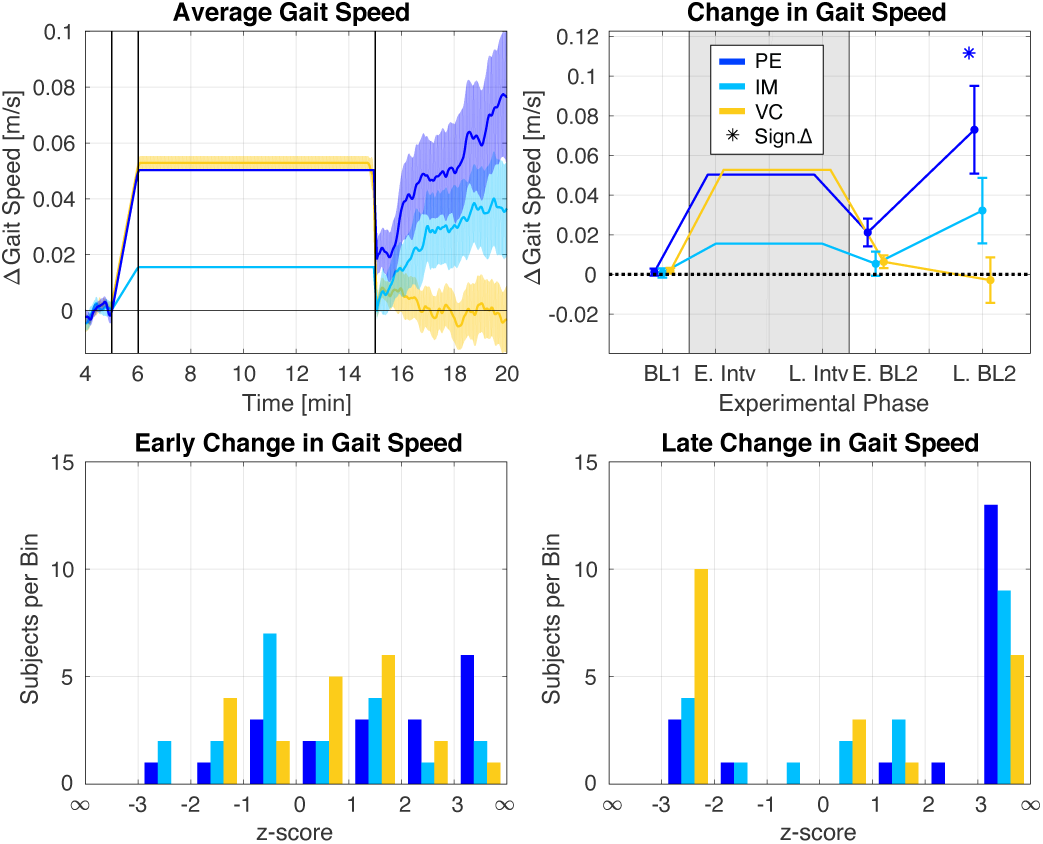
Within subject change in gait speed broken down by group. Representation of within subject change is provided for easier graphical display of the effects of training. Statistical analysis was performed on raw data. Top Left: Group average change in GS across the experimental protocol, resampled in time. Top Right: Mean and standard error of group average change in GS. Asterisks denote significant change from BL1 from post-hoc Tukey HSD analysis. Bottom: Histogram from responder analysis of Early (right) and Late (left) after effects following training.

**Fig. 6.**
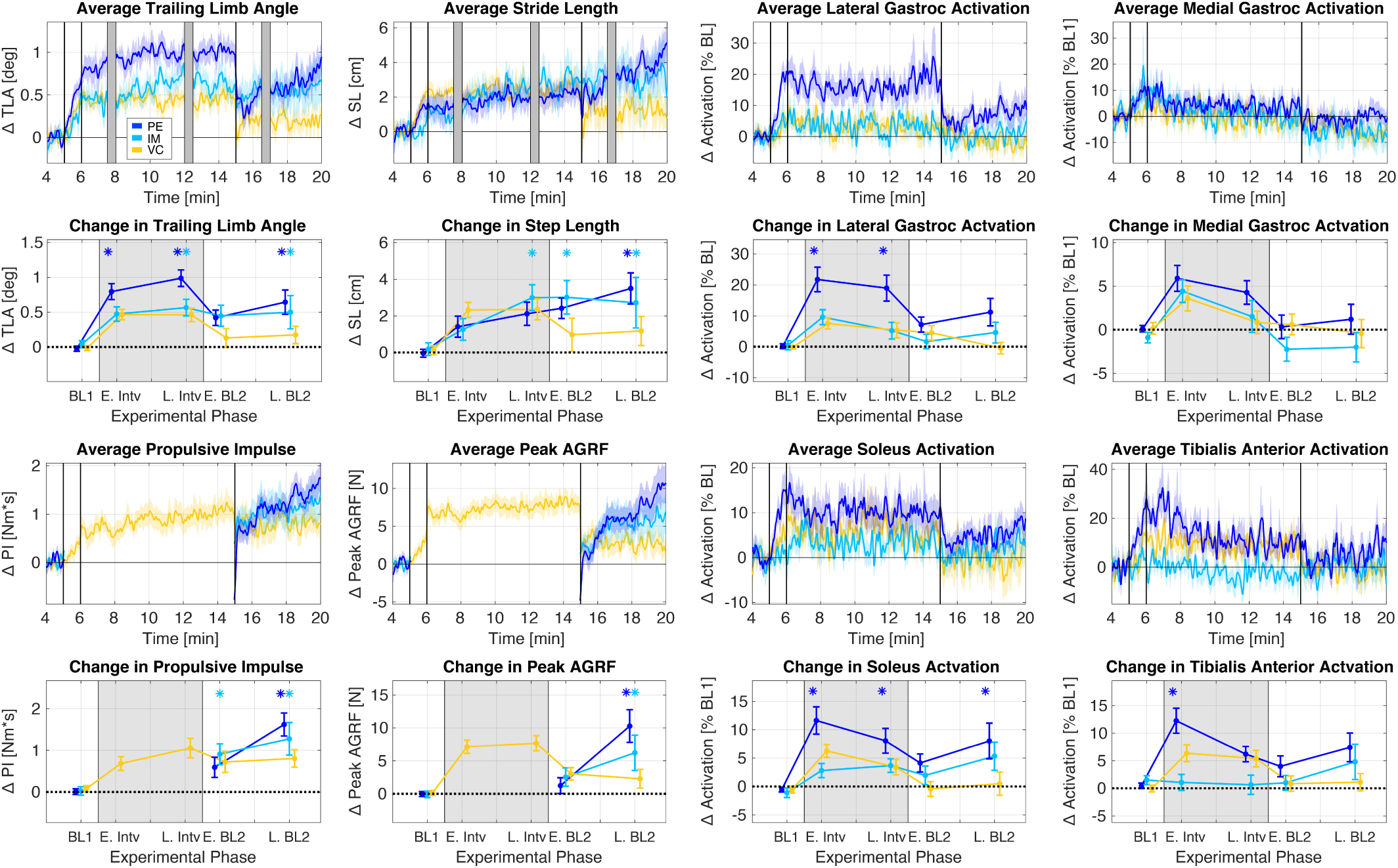
Within subject change in all gait parameters measured broken down by group. Rows 1 and 3: Group average change in gait parameters across the experimental protocol, resampled in time. Representation of within subject change is provided for easier graphical display of the effects of training. Statistical analysis was performed on raw data. Rows 2 and 4: Mean and standard error of group average change in gait parameters from our mixed model analysis. Asterisks denote significant change from BL1 from post-hoc Tukey HSD analysis.

### B. Walking Kinematics

#### 1) Trailing Limb Angle

The model returned a significant fixed effect of experimental phase, however the interaction between training type and experimental phase failed to reach significance (Tbl. I). No significant effect of group was detected. Model parameter estimates showed that across groups there was a significant increase in all experimental phases compared to BL1 (Early Intv.: 0.57 ± 0.08 deg, *p* < 0.001; Late. Intv.: 0.67 ± 0.08 deg, *p* < 0.001; Early BL2: 0.33 ± 0.08 deg, *p* < 0.001; Late BL2: 0.43 ± 0.08 deg, *p* < 0.001). Post-hoc Tukey HSD testing revealed a significant increase in trailing limb angle in Early Intv., Late Intv., and Late BL2 compared to BL1 in the PE group (0.80 ± 0.14 deg, *p* < 0.001; 0.99 ± 0.14 deg, *p* < 0.001; 0.64 ± 0.14 deg, *p* < 0.001, respectively), and in Late Intv and Late BL2 compared to BL1 in the IM group (0.57 ± 0.14 deg, *p* = 0.006; 0.50 ± 0.14 deg, *p* = 0.029, respectively). No significant change was detected in the VC group.

Responder analysis (Tbl. II, Fig. 7) showed a greater number of positive responders and larger median effects in the PE and IM groups compared to VC in BL2 (**PE:** Early: 16 (1.27 [0.13 2.80]), Late: 14 (2.24 [0.32 5.03]); **IM:** Early: 14 (1.42 [0.06 3.41]), Late: 15 (1.14 [0.05 5.08]); **VC:** Early: 11 (1.08 [0.16 2.24]), Late: 12 (1.00 [0.04 2.30])).

**Fig. 7.**
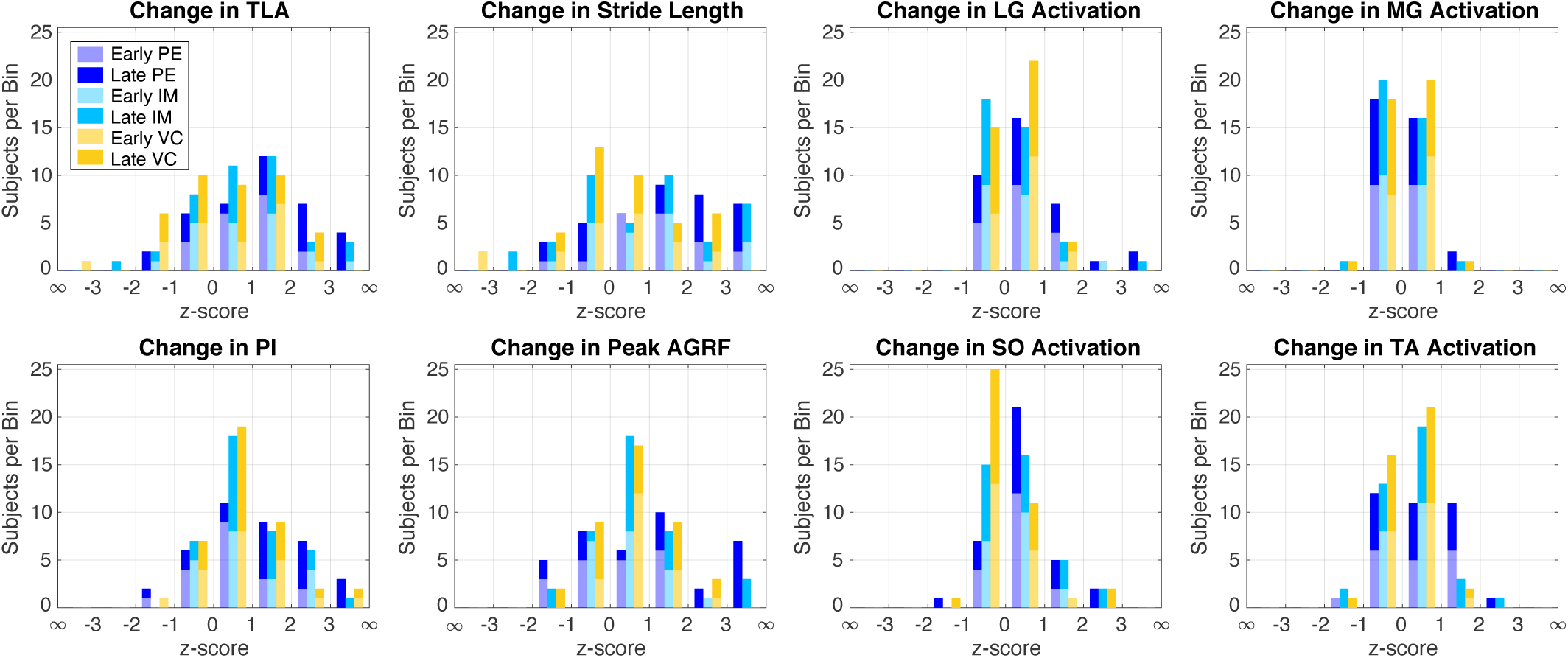
Histogram of responder analysis for early (strides 4-23) and late (final 20 strides) after effects following training for all gait parameters.

#### 2) Stride Length

The model returned a significant fixed effect of experimental phase, however the interaction between training type and experimental phase failed to reach significance (Tbl. I). No significant effect of group was detected. Model parameter estimates showed that across groups there was a significant increase in all experimental phases compared to BL1 (Early Intv.: 1.63 ± 0.46 cm, *p* = 0.004; Late Intv: 2.49 ± 0.46 cm, *p* < 0.001; Early BL2: 2.13 ± 0.46 cm, *p* < 0.001; Late BL2: 2.47 ± 0.46 cm, *p* < 0.001). Post-hoc Tukey HSD testing revealed a significant increase in stride length in Late BL2 compared to BL1 in the PE group (3.50 ± 0.81 cm, *p* = 0.002), and in Late Intv., Early BL2 and Late BL2 compared to BL1 in the IM group (3.00 ± 0.79 cm, *p* = 0.015; 3.02 ± 0.79 cm, *p* = 0.013; 2.72 ± 0.79 cm, *p* = 0.046, respectively). No significant change was detected in the VC group.

Responder analysis (Tbl. II, Fig. 7) showed a greater number of positive responders and larger median effects in the PE and IM groups compared to VC in BL2 (**PE:** Early: 17 (1.23 [0.24 3.36]), Late: 13 (2.78 [1.13 5.37]); **IM:** Early: 14 (1.38 [0.05 5.09]), Late: 11 (2.68 [0.27 9.53]); **VC:** Early: 11 (0.72 [0.07 2.47]), Late: 10 (1.16 [0.46 2.73])).

### C. Walking Kinetics

#### 1) Propulsive Impulse

The model returned a significant fixed effect of experimental phase, however the interaction between training type and experimental phase failed to meet our significance threshold (Tbl. I). No significant effect of group was detected. Model parameter estimates showed that across groups there was a significant increase in Early and Late BL2 over BL1 (Early BL2: 0.91 ± 0.26 Nm · s, *p* < 0.001; Late BL2: 1.28 ± 0.26 Nm · s, *p* = 0.001). Post-hoc Tukey HSD testing revealed a significant increase in propulsive impulse in Late BL2 compared to both BL1 and Early BL2 in the PE group (1.63 ± 0.27 Nm · s, *p* < 0.001; 1.03 ± 0.27 Nm · s, *p* = 0.005, respectively), and a significant increase in Early and Late BL2 compared to BL1 in the IM group (0.91 ± 0.26 Nm · s, *p* = 0.018; 1.28 ± 0.26 Nm · s, *p* < 0.001, respectively). No significant change was detected in the VC group.

Responder analysis (Tbl. II, Fig. 7) showed a similar number of positive responders between groups in BL2, but a greater median effect size for positive responders in the PE group in Late BL2 (**PE:** Early: 14 (0.69 [0.06 2.36]), Late: 16 (2.06 [0.31 4.46]); **IM:** Early: 15 (0.98 [0.07 2.99]), Late: 18 (0.83 [0.01 5.48] z-score); **VC:** Early: 15 0.83 [0.10 3.72]), Late: 17 (0.75 [0.23 3.34])).

#### 2) Peak AGRF

The model returned a significant fixed effect of experimental phase, and a significant interaction between training type and experimental phase (Tbl. I). No significant effect of group was detected. Model parameter estimates showed that across groups there was a significant increase in Early and Late BL2 over BL1 (2.27 ± 1.04 N, *p* = 0.031; 6.27 ± 1.04 N, *p* < 0.001), and in Late BL2 over Early BL2 (3.99 ± 1.04 N, *p* < 0.001). The interaction was driven by a significantly greater increase in Late BL2 from Early BL2 in the PE group compared to the IM and VC groups (**PE**: 9.05 ± 1.83 N; **IM**: 3.68 ± 1.79 N; **VC**: 0.75 ± 1.79 N; **PE vs IM:** *p* < 0.038; **PE vs VC:** *p* = 0.012), and a greater increase in Late BL2 from BL1 in the PE group compared to the VC group (**PE:** 10.30 ± 1.83 N; **VC:** 2.28 ± 1.79 N, *p* < 0.001). Post-hoc Tukey HSD testing revealed a significant increase in peak AGRF in Late BL2 compared to both BL1 and Early BL2 in the PE group (10.30 ± 1.83, *p* < 0.001; 9.05 ± 1.83,*p* < 0.001, respectively), and a significant increase in Late BL2 compared to BL1 in the IM group (6.22 ± 1.79, *p* = 0.020). No significant change was detected in the VC group.

Responder analysis (Tbl. II, Fig. 7) showed a greater number of positive responders in the VC group in Early BL2, but a greater number of positive responders in the PE and IM groups in Late BL2. Positive responders in the PE group had the largest median effect size across BL2 (**PE:** Early: 11 (1.13 [0.00 1.62]), Late: 14 (2.98 [0.66 5.35]); **IM:** Early: 13 (0.92 [0.04 2.34]), Late: 17 (0.72 [0.12 5.65]); **VC:** Early: 17 (0.75 [0.06 2.29]), Late: 12 (1.23 [0.06 2.55])).

### D. Muscle Activation

#### 1) Lateral Gastrocnemius

The model returned a significant fixed effect of experimental phase, and a significant interaction between training type and experimental phase (Tbl. I). No significant effect of group was detected. Model parameter estimates showed that across groups there was a significant increase in Early Intv., Late Intv., and Late BL2 over BL1 (Early Intv.: 12.98 ± 1.81%, *p* < 0.001; Late Intv.: 9.84 ± 1.81%, *p* < 0.001; Late BL2: 5.11 ± 1.81%, *p* = 0.042). The interaction was driven by a significantly greater increase in the PE group compared to the IM and VC groups in Early Intv. from BL1 (**PE:** 21.78 ± 3.22%; **IM**: 9.61 ± 3.13%; **VC:** 7.56 ± 3.05%; **PE vs IM:** *p* = 0.007; **PE vs VC:** *p* = 0.002), and in Late Intv. from BL1 (**PE:** 18.98 ± 3.22%; **IM:** 5.21 ± 3.13%; **VC:** 5.31 ± 3.05%; **PE vs IM:** *p* = 0.002; **PE vs VC:** *p* = 0.002), as well as a significantly greater increase in the PE group compared to VC group in Late BL2 over Early BL2 (**PE:** 4.04 ± 3.22%, **VC:** -4.98 ± 3.05%; *p* = 0.043) and in Late BL2 over BL1 (**PE:** 11.22 ± 3.22%, **VC:** -0.48 ± 3.05%; *p* = 0.009). Post-hoc Tukey HSD testing revealed a significant increase in activation in Early and Late Intv. compared to BL1 in the PE group (21.78 ± 3.22%, *p* < 0.001; 18.98 ± 3.22%, *p* < 0.001, respectively). No significant change in activation was detected in the IM or the VC group.

Responder analysis (Tbl. II, Fig. 7) showed a similar number of positive responders in all groups, but a larger median effect size in the PE group compared to the IM and VC group in Early BL2, and larger median effect both in PE and IM compared to VC in Late BL2 (**PE:** Early: 13 (0.74 [0.10 1.70]), Late: 13 (0.93 [0.02 3.75]); **IM:** Early: 10 (0.39 [0.09 2.23]), Late: 10 (0.55 [0.20 3.03]); **VC:** Early: 13 (0.43 [0.02 1.69]), Late: 10 (0.20 [0.01 1.25])).

#### 2) Medial Gastrocnemius

The model returned a significant effect of experimental phase. No significant effect of group or interaction was detected (Tbl. I). Model parameter estimates showed that across groups Early Intv. was significant greater than BL1 (mean ± s.e.m.: 4.65 ± 0.93%, *p* < 0.001).

In line with the group analysis, responder analysis (Tbl. II, Fig. 7) showed small, equivocal changes in MG activation in BL2 (**PE:** Early: 9 (0.36 [0.03 0.89]), Late: 9 (0.30 [0.01 1.65]); **IM:** Early: 9 (0.18 [0.01 0.82]), Late: 8 (0.22 [0.02 1.15]); **VC:** Early: 12 (0.22 [0.01 0.72]), Late: 9 (0.36 [0.03 1.02])).

#### 3) Soleus

The model returned a significant fixed effect of experimental phase, and a significant interaction between training type and experimental phase (Tbl. I). No significant effect of group was detected. Model parameter estimates showed that across groups there was a significant increase in Early Intv., Late Intv., and Late BL2 over BL1 (Early Intv.: 6.90 ± 1.13%, *p* < 0.001; Late Intv.: 5.04 ± 1.13%, *p* < 0.001; Late BL2: 4.61 ± 1.13%, *p* < 0.001). The interaction was driven by a significantly greater increase in Early Intv over BL1 in the PE group compared to the IM and VC groups (**PE:** 11.64 ± 2.0; **IM:** 2.8 ± 2.0%; **VC:** 6.2 ± 1.9%; **PE vs IM:** *p* = 0.002; **PE vs VC:** *p* = 0.05), and a significantly greater increase in Late BL2 over BL1 in the PE group compared to the VC group (**PE:** 8.0 ± 2.0; **VC:** -0.5 ± 1.9%, *p* = 0.007). Post-hoc Tukey HSD testing revealed a significant increase in activation in Early Intv., Late Intv., and Late BL2 compared to BL1 in the PE group (Early Intv.: 11.64 ± 2.0%, *p* < 0.001; Late Intv.: 8.1 ± 2.0%, *p* = 0.007; Late BL2: 8.0 ± 2.0%, *p* = 0.008). No significant change in activation was detected in the IM or the VC group.

Responder analysis (Tbl. II, Fig. 7) showed a greater number of positive responders in the PE and IM groups compared to the VC group in BL2 (**PE:** Early: 14 (0.50 [0.06 1.38]), Late: 14 (0.63 [0.10 2.95]); **IM:** Early: 12 (0.30 [0.09 1.46]), Late: 11 (0.88 [0.15 2.82]); **VC:** Early: 6 (0.59 [0.05 1.20]), Late: 7 (0.34 [0.05 2.17])).

#### 4) Tibialis Anterior

The model returned a significant fixed effect of experimental phase, and a significant interaction between training type and experimental phase (Tbl. I). No significant effect of group was detected. Model parameter estimates showed that across groups there was a significant increase in Early Intv., Late Intv., and Late BL2 over BL1 (Early Intv.: 6.57 ± 1.23%, *p* < 0.001; Late Intv.: 4.08 ± 1.23%, *p* = 0.01; Late BL2: 4.44 ± 1.23%, *p* = 0.004). The interaction was driven by a significantly greater increase in Early Intv. over BL1 in the PE group compared to the IM group (**PE:** 12.3 ± 2.2%; **IM:** 1.1 ± 2.1%, *p* < 0.001), and a significantly greater increase in Late BL2 over BL1 in the PE group compared to the VC group (**PE:** 7.4 ± 2.2%; **VC:** 1.1 ± 2.1%; *p* = 0.038). Post-hoc Tukey HSD testing revealed a significant increase in activation in Early Intv compared to BL1 in the PE group (12.3 ± 2.2, *p* < 0.001). No significant change in activation the IM or the VC group.

Responder analysis (Tbl. II, Fig. 7) showed a similar number of positive responders in all groups, but a larger median effect in the PE group compared to the IM and VC group in Early and Late BL2 (**PE:** Early: 11 (1.00 [0.27 1.21]), Late: 12 (0.97 [0.08 2.82]); **IM:** Early: 11 (0.26 [0.10 0.93]), Late: 12 (0.45 [0.12 2.22]); **VC:** Early: 12 (0.31 [0.11 1.24]), Late: 11 (0.27 [0.04 1.56])).

## IV. Discussion

We presented a novel paradigm used to train two components of propulsion during walking (push-off posture and ankle moment) to increase gait speed, based on the application of belt accelerations to the trailing limb during the double support phase of gait. In our protocol, we exposed two groups of subjects to belt accelerations at different magnitudes, Perceptible (7m/s^2^) and Imperceptible (2m/s^2^), and included a Velocity control group that was matched to the change in speed experienced in the Perceptible training group.

At the group level, the PE training group had the largest significant increase in gait speed following training (Late BL2: 0.073 ± 0.013 m/s, *p* = 0.005), and the greatest percentage of subjects with positive increases in gait speed (79%). The IM group exhibited a smaller effect (Late BL2: 0.032 ± 0.013 m/s, *p* = 0.25), and had fewer positive responders by the end of training (70%). The VC group had no significant change in velocity (Late BL2: -0.003 ± 0.013 m/s), and a 50-50 split between positive and negative responders.

We hypothesized that during training subjects would need to push off harder and that accelerations would increase trailing limb angle. Analysis of gait kinematics show that significant increases in TLA were seen in our PE and IM groups during training (PE: 0.99 ± 0.14 deg, p < 0.001; IM: 0.57 ± 0.14 deg, *p* = 0.006). Although the VC group was velocity matched to the PE group, changes in gait kinematics were smaller in magnitude and non significant, which suggests that accelerations were more effective in stimulating increases in TLA compared to increases in velocity alone (VC TLA: 0.46 ± 0.14 deg). Following training, the VC group returned to their baseline TLA, while both acceleration training conditions maintained the trained increases in TLA (PE: 0.64 ± 0.14 deg, p < 0.001; IM: 0.50 ± 0.14 deg, *p* = 0.03). Stride length, another means of quantifying postural change, showed similar results (PE Intv: 2.12 ± 0.81 cm, BL2: 3.50 ± 0.81 cm, *p* = 0.002; IM Intv: 3.00 ± 0.79 cm, *p* = 0.015, BL2: 2.72 ± 0.79 cm, *p* = 0.046; VC Intv: 2.36 ± 0.80 cm, BL2: 1.17 ± 0.80 cm).

During acceleration training, force-plate measurements of propulsion are contaminated by accelerations of the treadmill belt [21]. In the VC group, in which this is not the case, significant increases in PI and peak AGRF were measured (PI: 1.06 ± 0.24 Nm*s; peak AGRF: 7.68 ± 1.63 N), as expected. During training, EMG measures of the lateral gastrocnemius and soleus plantar flexor muscles showed that increases in activation in the PE group were significantly greater than in the VC group. From these measures, we can infer that greater ankle moment was required to respond to belt accelerations during push-off compared to a simple increase in belt velocity experience in the VC group, as expected. Following training, the increases in plantar-flexor muscle activity were sustained in the PE group, as were increases in propulsive impulse and peak AGRF. Changes in activation in the Tibialis Anterior, a dorsiflexor muscle, are in line with previous research that show increased activation with gait speed for velocities greater than self-selected walking speed [5]. The results for each training group are in line with their respective changes in velocity.

To investigate type of learning driving potential gait modifications, we compared gait parameters measured in early and late training, and early and late post-training. If adaptation drives gait modifications, we expect to see significant change in behavior over training as subjects adapt to the perturbation. Following training, we expect to see after effects in the opposite direction of change during training. Moreover, we would expect to see different timescales of retention in the PE and IM training groups, with large, immediate after effects in the PE training group that decay quickly (within 1 minute), and smaller immediate after effects that decay slowly in the IM training group (within 5 minutes). Instead, in UDL we expect to see no significant change in behavior between early and late training. Because after effects in UDL reflect the trained repeated movement, we expect to see greater after effects in the PE training group, as it induces a greater gait modifications during training, and smaller effects in the IM training group. We do not expect to see any differences in the timescale of retention in after effects driven by UDL between PE and IM training groups, but expect them to last for more than 5 minutes [15]–[17].

In all training groups, change measured between Early and Late Intervention was either not significant, or decreased. In both groups exposed to belt accelerations, after effects were in the same direction as the effect of training, scaled with the magnitidue of change during training, and persisted across the baseline 2 walking period. These results are inconsistent with adaptation, in both timescale and direction. Adaptation after effects generally decay more rapidly, and occur in the opposite direction of adaptation during training, suggesting that after effects should be less than BL1 immediately following training, which was not observed in any parameter tested. Instead, it appears that our protocol stimulates UDL, as all after effects are seen to occur the same direction as the change during training and persist for the duration of the baseline 2 experimental phase. The resultant increase over the course of BL2 is not consistent with traditional washout of UDL in a null condition. However, the nature of the UDTC allows for modifications in gait to be maintained, and may reinforce the trained behavior, allowing for increases in BL2 that approach the magnitude of the gait modifications applied during training [23]. Moreover, biomechanical changes in gait have been shown to precede changes in gait speed, and it is not a simple one-to-one relationship between biomechanics and gait-speed outcome as is commonly studied in the UDL literature [17], [22], [23]

The number of non-responders in both training groups is also in line with UDL given the short duration of our training protocol (10 minutes). Studies of repetitive movements have shown that the number of movement repetitions influences whether or not UDL occurs [17], [23]. In repetitive thumb movements, 10 minutes of training at a 1 Hz movement frequency resulted in only 33% of subjects showing significant UDL, while 15 minutes increased this percentage to 50%. 100% of subjects showed significant UDL after 30 minutes of training [17]. Step frequency in our training protocol was roughly 2 Hz, putting the number of movement repetitions in our protocol roughly between the 15-30 minute training groups in the Classen study. As such, the percentage of expected responders should be between 50 and 100%, which is in agreement with the 70-79% response rate achieved in our study. Future work will look at increasing training time to determine if increased repetition in our training protocol increases the number of positive responders.

Our responder analysis further elucidated differences in group behavior that highlight which gait parameters may have contributed most to differences in the final gait speed measured between groups. In the acceleration training groups, a greater number of subjects had early increases in TLA (PE: 16, IM: 14, VC: 11), SL (PE: 17, IM: 14, VC: 11) and Soleus activation (PE: 14, IM: 11, VC: 7) immediately following training that was sustained across BL2. While a similar number of subjects increased Lateral Gastroc activation (PE: 13, IM: 10, VC: 13) across groups immediately following training, increases in the PE training group were almost twice as large as those measured in the other two groups. Evaluation of early propulsion shows no significant differences between groups, and a slightly larger increase in peak AGRF in the VC training group. From these results, it seems that the learned changes in gait kinematics paired with sustained change in the neuromotor commands sent to plantar-flexor muscles contributed to the larger increases in gait speed and propulsion measured at the end of training in our accelerating training conditions compared to velocity control.

The end magnitude of change in gait speed in the PE training group was 0.073 m/s. Current gait training strategies for stroke survivors achieve an average change in gait speed of 0.06 m/s [24]. Minimal clinically significant change in gait speed is 0.16 m/s, and can take up to 36 sessions of training to achieve [22]. While the effects in our study in healthy subjects do not directly translate to a patient population, the magnitude of our effects after just one training session are encouraging for future research in a patient population, especially considering that healthy subjects are subject to ceiling effects as their baseline is assumed to be roughly optimal. Moreover, previous studies have shown that training increases in muscle strength alone do not translate to functional increases in ankle moment during walking, and call for rehabilitation methods that target propulsive deficits in the context of walking [1]. Our paradigm has the potential to improve locomotor patterns through training favorable biomechanical changes in both posture and ankle plantar-flexors during walking, with potential to provide greater translation to habitual speed walking in the community setting than muscle strengthening alone, or other gait therapies that utilize fixed speed treadmills [1], [3], [18], [24].

## V. Conclusion

Overall, our paradigm significantly increased walking speed in 79% subjects in the Perceptible group and 70% in the Imperceptible group. Changes in gait speed were greater in the Perceptible training group. The Imperceptible training group had smaller but qualitatively similar effects in the same direction as the Perceptible training group, suggesting that the magnitude of after effects scales with the magnitude of belt acceleration applied during training. Changes in gait speed measured after training were driven by both changes in ankle moment— measured by immediate and sustained increases in activation in plantar-flexor muscles, and increases in PI and Peak AGRF over baseline 2—and by push off posture, measured as the sustained increases in TLA and SL in both acceleration training groups. Our results indicate that the Perceptible intervention is a better candidate for continued research in training increased gait speed, as subjects in this group had more consistent after-effects, larger increases in all gait parameters tested, and sustained change in walking behavior following intervention. Future work will focus on identifying the factors that contribute to variable response of subjects to training, including duration of training, and the extension of our work in healthy individuals to a patient population, or to healthy elderly adults.

## VI. Acknowledgments

We acknowledge support from the NSF grant no. 1638007, and from startup funds by the University of Delaware. We thank Andrew Borowski, Rachel Marbaker and Nicole Ray for their collaboration on the treadmill controllers.

---

**Figure 1:**
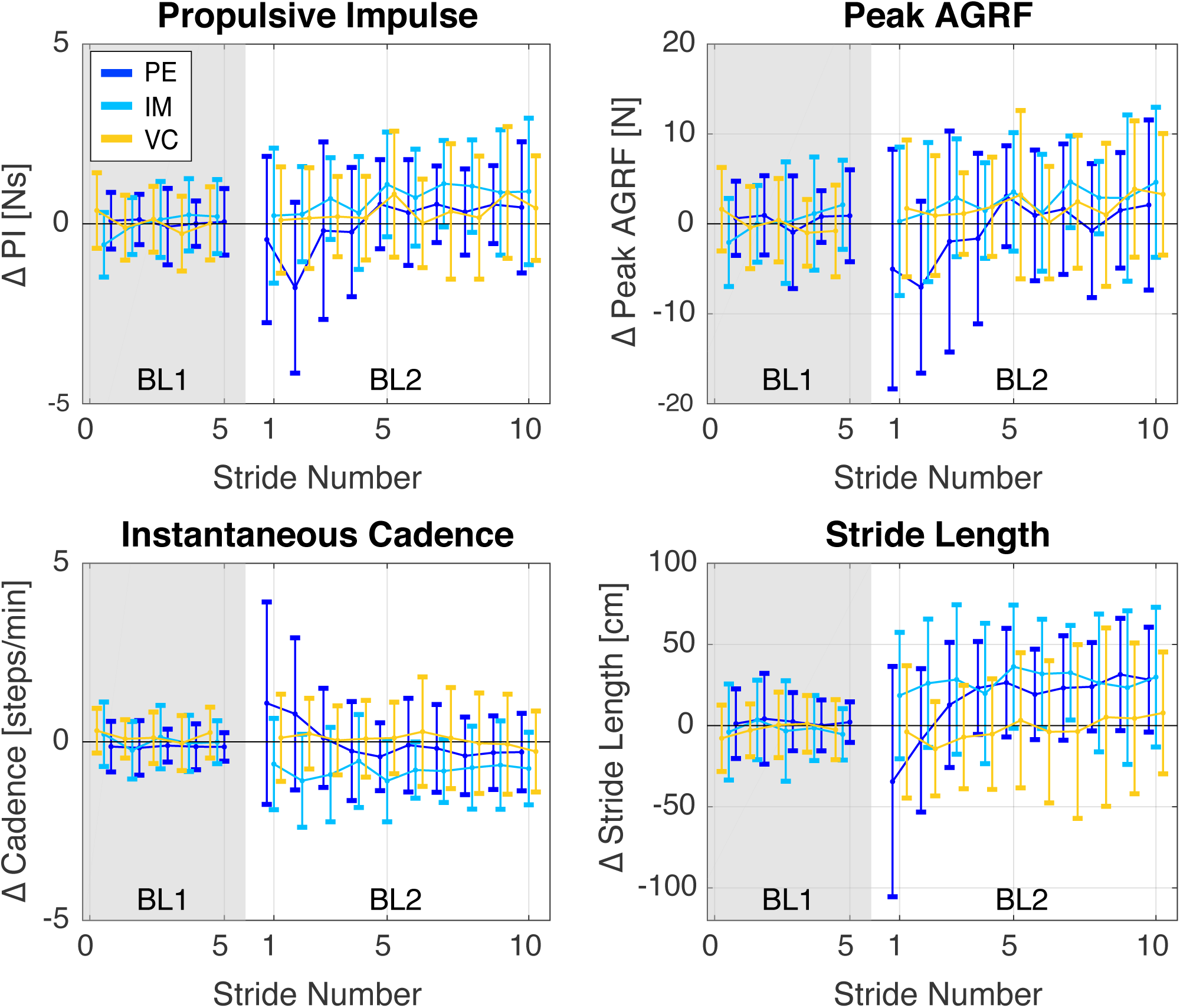
Gait parameters measured in the last 5 strides of BL1 are shown compared to the first 10 strides of BL2. Data shows a transient period in the extent of roughly four stides in BL2 in the Perceptible training group only. Behavior is consistent with stutter steps, in which subjects take rapid, shorter steps as they adjust to the change in belt behavior, evidenced by the measured decreases in stride length, increases in instantaneous cadence, as well as the decreased propulsive forces. A transient response when belt behavior is changed during active walking has been previously reported [20]. There was no transient behavior measured in plantar-flexor muscle activity or TLA immediately following training. To be consistent across parameters, and avoid undue influence by transient behavior, we excluded transient strides 1-4 from our analysis and defined Early BL2 as strides 5-24 for all gait parameters in our data analysis.

